## Supplementary material and methods for "Evolutionary unique *N*-glycan-dependent protein quality control system plays pivotal roles in cellular fitness and extracellular vesicle transport in *Cryptococcus neoformans*"

**Supplementary File 1**

**Supplementary Materials and Methods**

**Construction of *C. neoformans* *ugg1*Δ and *UGG1* complementation strains**

The *UGG1* (CNAG_03648) deletion mutant strain was constructed by amplifying the DNA fragments containing the 5′- or 3′-flanking regions of CNAG_03648 ORF using PCR, in which genomic DNA from the H99 strain and the following primer sets: 03648p_Fw/03648p_Rv and 03648t_Fw/03648t_Rv, were used (Table S1C). The 5′- and 3′-regions of the selectable marker nourseothricin acetyltransferase (NAT) were amplified using the primer sets M13Fe/NSL-2 and M13Re/NSR-2 and pNAT-STM#159 as the template. *UGG1–NAT* fusion products of 5′- and 3′-flanking regions were generated using overlap PCR with the primer sets M13Fe/03648p_Rv and 03648t_F/NSR-2, respectively. The 5′- and 3′-fragments of the *UGG1* disruption cassette were introduced into *C. neoformans* serotype A strain H99 through biolistic transformation. The transformants were selected on YPD_NAT_, and gene disruption was screened using PCR. The complemented strain was generated by amplifying the DNA fragment containing *UGG1* using PCR, followed by subcloning into pJAFS1 containing the G418 resistance marker via In-Fusion cloning (Takara Bio Inc.). The resulting vector was excised at the unique EcoRV site and reintegrated into the native locus of the *ugg1*Δ strain through biolistic transformation. The transformants were selected on YPD_G418_, and gene disruption was screened using PCR.

**Construction of *C. neoformans* *mns1*Δ, *mns101*Δ, *mns1*Δ*101*Δ, and complementation strains**

The *MNSA* (CNAG_02081) and *MNS101* (CNAG_03240) deletion mutant strains were constructed by amplifying the DNA fragments containing the 5′- or 3′-flanking regions of CNAG_02081 and CNAG_03240 ORFs using PCR with the genomic DNA from the H99 strain and the primer sets: 02081_L1/02081_R1, 02081_L2/02081_R2 and 03240_L1/03240_R1, 03240_L2/03240_R2, respectively (Table S1C). The 5′- and 3′-regions of the selection marker Hygromycin B (HyB) were amplified using the primer sets M13Fe/B5752 and M13Re/B5751 and using pJAF_HyG as the template. *MNS1-HyB* fusion products of 5′- and 3′-flanking regions were generated through overlap PCR using the primer sets M13Fe/02081_R1 and 02081_L2/NSR-2, respectively. *MNS101-NAT* fusion products of 5′- and 3′-flanking regions were generated through overlap PCR and the primer sets M13Fe/03240_R1 and 03240_L2/NSR-2, respectively. The 5′- and 3′-fragments of the *MNS1* and *MNS101* disruption cassettes were introduced into *C. neoformans* serotype A strain H99 through biolistic transformation. The transformants were selected on YPD_HyB_ or YPD_NAT_, respectively, and gene disruption was screened using PCR. The mutant strain lacking both *MNS1* and *MNS101* (*mns1*Δ*101*Δ) were generated by introducing the *MNS1* disruption cassette into the *mns101*Δ deletion strain. Each complemented strain was generated by amplifying the DNA fragment containing *MNS1* or *MNS101* using PCR, followed by subcloning into pJAFS1 containing the G418 resistance marker via In-Fusion cloning (Takara Bio Inc.). The resulting vector was excised at the unique BstBI site and reintegrated into the native locus of the *mns1*Δ, *mns101*Δ, or *mns1*Δ*101*Δ strains through biolistic transformation. The transformants were selected on YPD_G418_, and gene disruption was screened using PCR.

**Construction of *C. neoformans* *mnl1*Δ, *mnl2*Δ, and *mnl1*Δ*mnl2*Δ and complementation strains**

The *MNL1* (CNAG_01987) and *MNL2* (CNAG_04498) deletion mutant strains were constructed by amplifying the DNA fragments of the 5′- or 3′-flanking regions of CNAG_01987 and CNAG_04498 ORFs by PCR using genomic DNA from the H99 strain and the primer sets 01987_L1/01987_R1, 01987_L2/01987_R2 and 04498_L1/04498_R1, 04498_L2/04498_R2, respectively (Table S1C). The *MNL1* disruption was performed by amplifying the 5′- and 3′-regions of the selectable marker nourseothricin acetyltransferase (NAT) with the primer sets M13Fe/NSL-2 and M13Re/NSR-2 using pNAT-STM#159 as a template. The *MNL1-NAT* fusion products of the 5′- and 3′-flanking regions were generated using overlap PCR with the primer sets M13Fe/01987_L2 and 01987_L1/NSR-2, respectively. The *MNL2* disruption was performed by amplifying the 5′- and 3′-regions of the selectable marker G418 with the primer sets M13Fe/B1887 and M13Re/B1886 using pJAFS1 as a template. The *MNL2-G418* fusion products of 5′- and 3′-flanking regions were generated using overlap PCR and the primer sets M13Fe/04498_R2 and 04498_F1/B1887, respectively. The 5′- and 3′-fragments of each *MNL1* and *MNL2* disruption cassettes were introduced into *C. neoformans* serotype A strain H99 through biolistic transformation. The transformants were selected on YPD_NAT_ or YPD_G418_, respectively, and gene disruption was screened using PCR. Mutant strain lacking both *MNL1* and *MNL2* (*mnl1*Δ*mnl2*Δ) were generated by introducing the *MNL2* disruption cassette into the *mnl1*Δ deletion strain. Each complemented strain was generated by amplifying the DNA fragment containing *MNL1* or *MNL2* using PCR and subcloned into pJAFS1 containing the G418 resistance marker. The transformants were selected on YPD_NAT_, and gene disruption was screened using PCR.

**Construction and localization analysis of *C. neoformans* GFP-Ugg1, Mns1-GFP and Mns101-GFP fusion proteins**

The GFP-UGG1 fusion vector expressing an N-terminal GFP fusion protein was generated by amplifying the *UGG1* and the *C. neoformans* codon-optimized GFP sequences using the primer sets Bv_Not1_ugg1_pro_F1/SP_03648pro_R1, SP_CnGFP_F2/UGG1ORF_5gly_CnGFP_R2 and UGG1ORF_5gly_CnGFP_F3/Bv_Mfe1_UGG1ORF_R3 with pJAFS1_UGG1Com and pWH091 as templates. The fused PCR product was subcloned into the pJAFS1_UGG1Com plasmid via In-Fusion cloning (Takara Bio Inc.) to generate the pJAFS1_SP_CnGFP_UGG1 plasmid. The resultant plasmid pJAFS1_SP_CnGFP_UGG1 was digested using EcoRV and biolistically transformed by integration into the CNAG_03648 locus of the *ugg1*Δ strain. The MNS1-GFP and MNS101-GFP fusion vectors expressing a C-terminal GFP fusion protein were generated by amplifying the *MNS1* and *MNS101* ORF sequences using the primer sets BamHI_MNS1 F1/MNS1_GFP R1 and BamHI_MNS101 F1/MNS101_GFP R1, respectively, and subcloned via In-Fusion cloning (Takara Bio Inc., Japan) into the pWH091 plasmid to generate the pWH091-MNS1 and pWH091-MNS101 plasmids. Both plasmids were digested with BstB1 and integrated into the CNAG_02081 and CNAG_03240 locus, respectively of the WT strain locus by biolistic transformation.

The yeast cells were cultured in YPD medium for 24 h and the obtained cell pellets were fixed in 4% paraformaldehyde [pH 7.0] for 10 min in rotation. The cells were washed twice with PBS and stained with 1 µM ER Tracker^TM^ red (Invitrogen, USA) for 20 min and 5 µg/ml of DAPI (4′,6-diamidino-2-phenylindole) for 10 min. The cells were washed twice with PBS and adjusted to an OD_600_ of 1.0 in ultra-pure water. Fluorescence was observed through an Eclipse Ti-E fluorescence microscope (Nikon, Japan) equipped with a Nikon DS-Qi2 camera and a Plan Apo VC 100X Oil DIC N2 (NA 1.4) lens. Images were processed on the NIS-elements imaging software (Nikon, Japan).

**HPLC and MALDI-TOF-based *N*-glycan structure analysis**

Cell wall mannoproteins (cwMPs) from *C. neoformans* were isolated and subjected to *N*-glycan structure analysis as described previously (Park et al., 2012; Thak et al., 2018). Briefly, *N*-glycans were released from purified cwMPs through PNGase F treatment (New England Biolabs, UK), followed by purification using a Carbograph Extract-Clean column. The purified *N*-glycans were labeled with 2-aminobenzoic acid (2-AA; Sigma-Aldrich, USA) and further purified using a Cyano Base cartridge (Agilent, USA). 2-AA-labeled *N*-glycans were analyzed using a Waters 2690 HPLC system equipped with a 2475 fluorescence detector that was set to excitation and emission wavelengths of 360 nm and 425 nm, respectively. Data were collected using the Empower 2 software (Waters). For MALDI-TOF analysis, neutral N-glycans were collected from HPLC fractionation and dried. The matrix solution was prepared as previously described (Thak et al., 2018) and mixed with the samples in equal volumes. The samples were spotted on an MSP 96 polished-steek target (Bruker Daltonics, Germany), and the crystalized samples were analyzed using a Microflex mass spectrometer (Bruker Daltonics, Germany) in a linear negative mode.

**Animal study and *in vitro* survival analysis**

Animal studies were conducted at the Chung-Ang University Animal Experiment Center. The study design was approved by the Ministry of Food and Drug Safety (MFDS, South Korea). Survival and fungal burden were assayed as described previously (Cheon et al., 2011). Briefly, eight mice (6-week-old female A/J Slc mice; Japan SLC) per strain were infected with 10^5^ cells via intranasal instillation. The mice were weighed and monitored once daily and euthanized after rapid 30% weight loss or identification of signs of morbidity. Kaplan–Meier survival curves were generated using Prism version 7 (GraphPad Software). For performing the fungal burden assay, the lungs of *C. neoformans*-infected mice were dissected on days 7 or 60. Half-organ portions of the excised lungs were homogenized, serially diluted, and plated onto YPD medium containing 100 μg/ml chloramphenicol (Sigma-Aldrich, USA). The other half-lung samples were fixed, sectioned, and stained with mucicarmine (Abcam, UK) for histopathological analysis. *C. neoformans* colonization was analyzed using a Zeiss Axioscope (A1) equipped with an AxioCam MRm digital camera.

Cell survival within macrophages was analyzed by opsonizing *C. neoformans* cells with 10 mg/ml of 18B7 antibody at 37 °C for 1 h. The macrophage-like J774A.1 (10^5^) cells were seeded onto 96-well plates in DMEM medium supplemented with 10% FBS and cultured at 37 °C in 5% CO_2_ for 18 h. The opsonized *C. neoformans* (10^5^) cells were co-incubated with activated macrophages at 37 °C in 5% CO_2_ for 1 h. The non-phagocytized yeast cells were removed by washing each well thrice with PBS. Then, DMEM medium supplemented with 10% FBS was added to each well, followed by culturing at 37 °C in 5% CO_2_ for 24 h. The macrophages were lysed in distilled water by vigorous pipetting, and the fungal cells were collected and serially diluted. Cryptococcal survival was assessed using two independent colony forming unit (CFU) assays.

**Subcellular fractionation and western blot analysis of Cda1**

Subcellular fractionation was performed as previously described (Thak et al., 2022) for localization analysis of Cda1. Briefly, the respective strains were inoculated at an initial OD_600_ of 0.5 in YPD medium and cultured at 30 °C for 16 h. Cell suspensions (OD_600_ = 30) were divided into two tubes to extract total cellular proteins and to fractionate soluble/insoluble cellular proteins. The cells were disrupted four times using glass beads (425–600 μm in diameter, Sigma) for 15 s at 5,000 rpm in a Precellys^®^ 24 Tissue Homogenizer (Bertin Technologies, France). Total proteins were extracted by adding 5X sample loading buffer (62.5 mM Tris-HCl pH 6.8, 2.5% SDS, 0.002% Bromophenol Blue, 5% β-mercaptoethanol, and 10% glycerol) and boiling for 10 min. Cell debris and glass beads were removed by centrifugation for 1 min at 16,000 ×*g*. The soluble protein fraction was obtained after centrifugation for 10 min at 16,000 ×*g*, and 5X sample loading buffer was added to the supernatant, followed by boiling for 10 min. The insoluble protein fraction was obtained by adding 1X sample loading buffer into the remaining pellets and boiling for 10 min. The obtained samples were adjusted to the same protein concentration and separated using SDS-PAGE. Cda1 expression was analyzed suing western blotting with an anti-Cda1 monoclonal antibody (Upadhya et al., 2018).

**RNA preparation, qRT-PCR, and RNA-sequencing**

Total RNA preparation, qRT-PCR, and RNA-sequencing were performed as previously described (Thak et al., 2022). Cells were inoculated in YPD medium at an OD_600_ of 0.15 and cultured until it reached the mid-logarithmic phase (OD_600_ of 0.5). Total RNA was extracted using the RNeasy Mini Kit (Qiagen, Germany) and single stranded cDNA was synthesized using the Superior Script III reverse transcriptase (Enzynomics, South Korea). Reverse transcriptase-polymerase chain reaction (RT-PCR) was performed using serially diluted cDNA with the gene-specific primer sets (Supplementary Table S1C) using Maxime PCR PreMix (i-Taq) (iNtRON Biotechnology, South Korea). Quantitative real-time PCR (qRT-PCR) was performed with the gene-specific primer sets (Supplementary Table S1, C) using a CFX96 Real-Time PCR detection system (Bio-Rad, USA). Normalized fold expression was calculated with the CFX manager software using the 2^−ΔΔCT^ method and with ACT1 or *GAPDH* as the reference genes. For RNA-sequencing, the NEBNext Ultra Ⅱ Directional RNA-Seq Kit (New England BioLabs, UK) was used to construct libraries. The mRNA was isolated using the Poly(A) RNA Selection Kit (Lexogen, Austria) and used for the cDNA synthesis. NovaSeq 6000 (Illumina) was used for high-throughput sequencing as paired-end 100 sequencing. Adapter and low-quality reads (<Q20) were removed using FASTX_Trimmer and BBMap. Trimmed reads were mapped to the reference genome using TopHat, and the RC (Read Count) data were processed based on FPKM+Geometric normalization method using EdgeR. Gene expression levels were quantified by evaluating the Fragments Per kb per Million (FPKM) reads values using Cufflinks. Data mining and graphic visualization were performed using ExDEGA (Ebiogen Inc., Korea).

**Sample preparation for proteomic analysis**

Proteins from extracellular vesicles (EVs) and whole-cell lysates (WCL) were prepared for proteomic analysis as follows. For EV proteome analysis, proteins were extracted by solubilizing purified EVs in a solution containing 8 M urea, 100 mM Tris (pH 7.5), and 5 mM tris(2-carboxyethyl)phosphine (TCEP) for 20 min at 23 °C. For WCL samples, cell pellets collected after EV secretion were resuspended in the same extraction buffer and disrupted using glass beads (425–600 µm, Sigma-Aldrich, USA) with a Precellys 24 Tissue Homogenizer (Bertin Technologies, France) at 5,000 rpm for four 15-second cycles. Following disruption, samples were centrifuged at 16,000 rpm for 5 minutes at 4 °C, and the supernatants were collected as WCL samples. For secretome and WCL proteome analysis, *Cryptococcus neoformans* cells were cultured in synthetic dextrose (SD) liquid medium for 24 h. Culture supernatants were collected after centrifugation and filtered through 0.22 μm syringe filters to remove cellular debris, yielding secretome samples. Cell pellets were resuspended in 8 M urea extraction buffer and disrupted using glass beads as described above to generate WCL samples.

Total protein concentrations were quantified using the Protein Assay Dye (Bio-Rad, USA). Protein digestion was performed with Protifi S-Trap™ mini spin columns (C02-mini-80, Protifi, USA) and trypsin gold (V5280, Promega, USA). Digested peptides were eluted with 50 mM tetraethylammonium bromide (TEAB; Thermo Fisher Scientific, USA), 0.2% formic acid, and 50% acetonitrile. The pooled peptides were dried, dissolved in 100 mM TEAB, and labeled using the TMTpro™ 16plex Label Reagent Set (Thermo Fisher Scientific, USA).
