## Supplementary figures for "Evolutionary unique *N*-glycan-dependent protein quality control system plays pivotal roles in cellular fitness and extracellular vesicle transport in *Cryptococcus neoformans*"

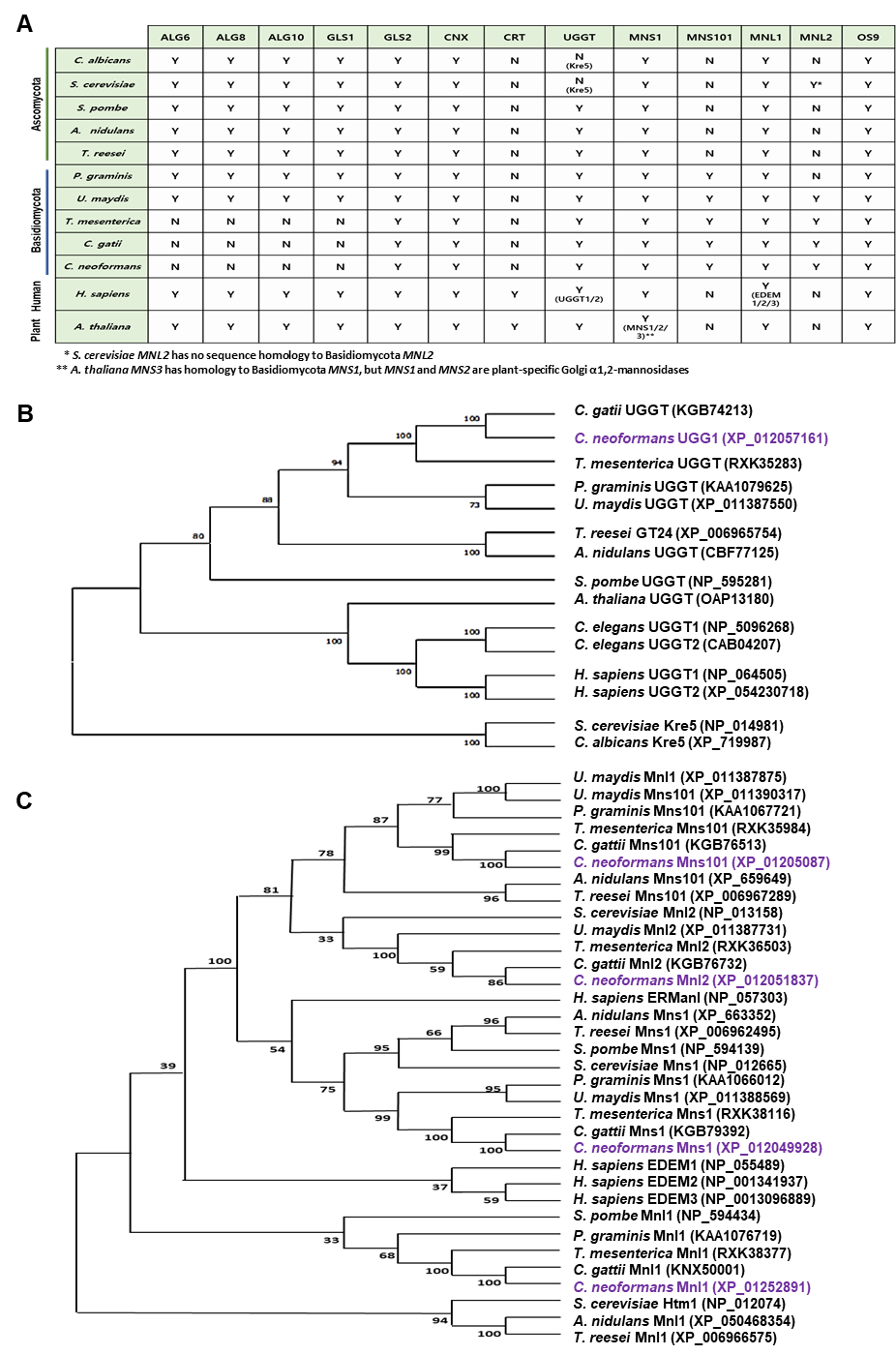
**Supplementary Figure S1.** **Evolutionary conservation and divergence of endoplasmic reticulum quality control (ERQC)-related gene homologs in eukaryotes.** (A) Distribution of ERQC homologs across various eukaryotic organisms. (B, C) Phylogenetic analysis of UDP-glucose:glycoprotein glucosyltransferase (UGGT)*,* α1,2-mannosidase I (Mns1) and α1,2-mannosidase-like proteins (Mnl1/Htm1/Mnl2) homologs in different fungi. The phylogenetic trees were constructed using the maximum likelihood method in the MEGA X software with bootstrap analysis performed using 100 random resampling.

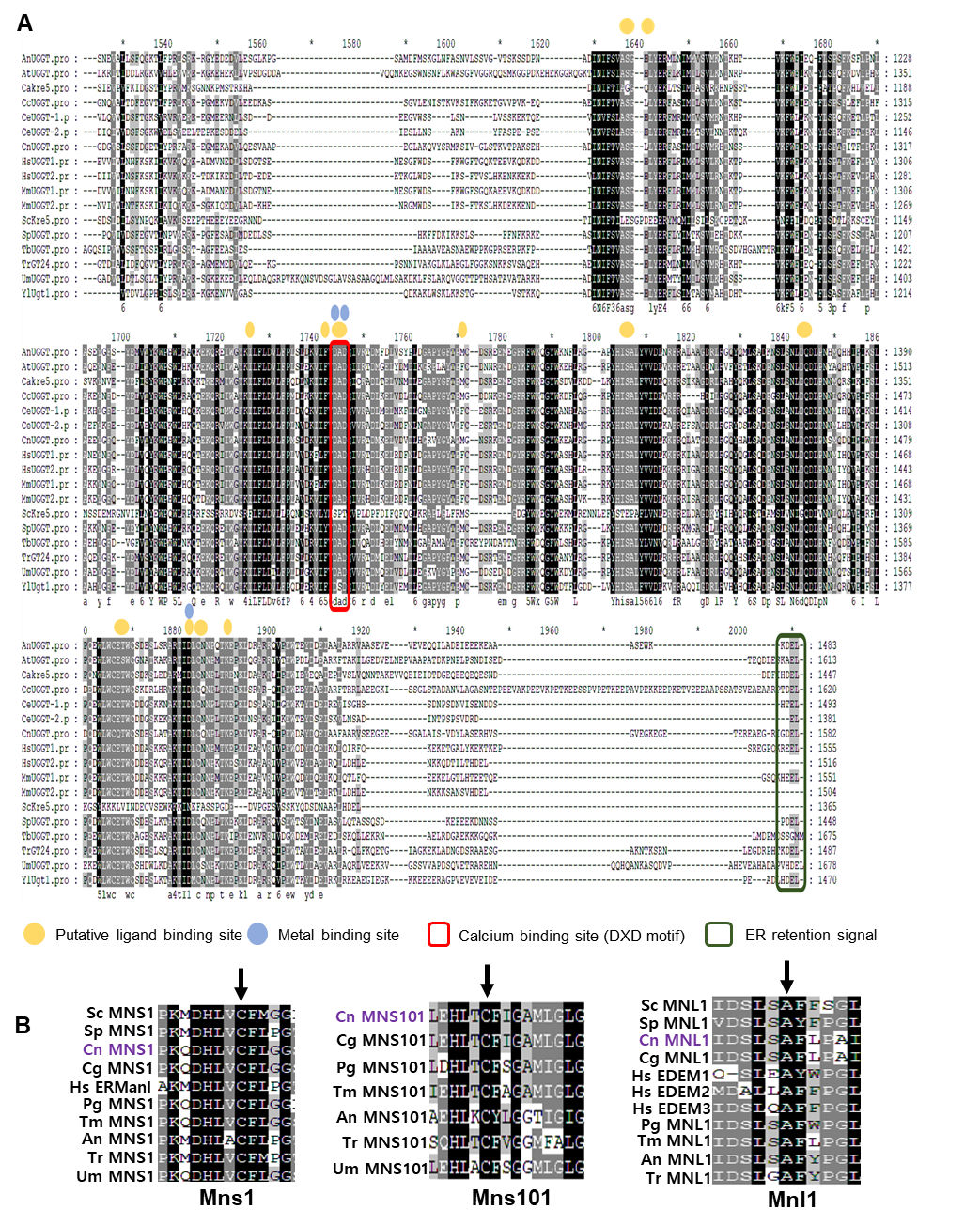

**Supplementary Figure S2. Domain analysis of putative *C. neoformans* ERQC-associated proteins.** (A) Multiple sequence alignment of UGGT homologs generated using Clustal W. Identical and conserved amino acids are indicated by black and gray shading, respectively. The calcium-binding site and HDEL-like ER retention signal are highlighted with boxes. (B) Multiple sequence alignment highlighting the conserved cysteine and alanine residues (indicated by the arrows) present in all putative Mns1 and Mnl1 homologs, which are essential for mannosidase activity. This alignment was generated using Clustal W 1.81 and shaded using the GeneDoc software.

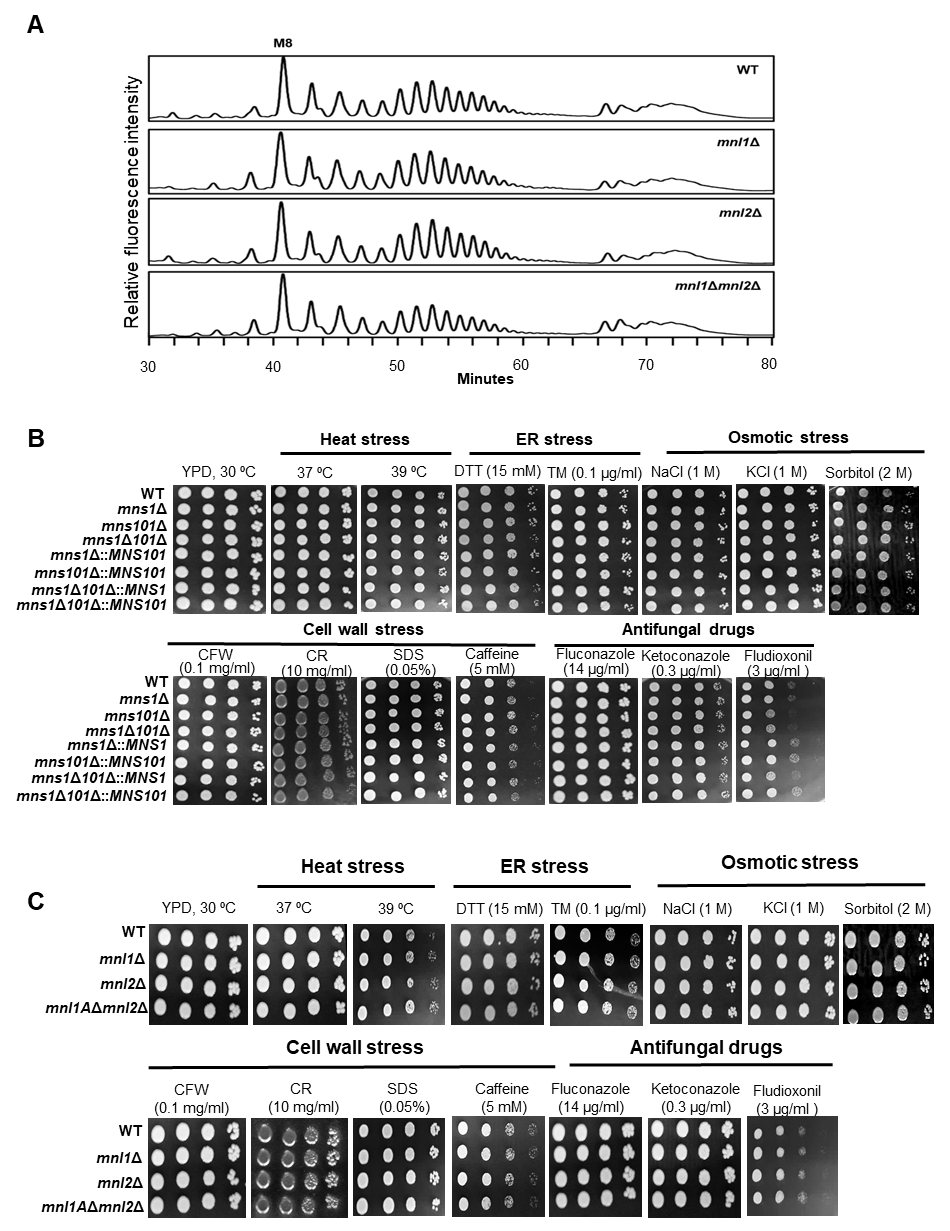

**Supplementary Figure S3. Phenotype characterization of *C. neoformans*** **α1,2*-*mannosidases.** (A) HPLC-based *N*-glycan profile of *C. neoformans mnl1*Δ, *mnl2*Δ, and *mnl1*Δ*mnl2*Δ mutants. (B) Spotting analysis of *C. neoformans mns1*Δ, *mns101*Δ, and *mns1*Δ*101*Δ cultivated under various stress conditions including heat stress (37 °C, 39 °C), ER stress (DTT: dithiothreitol, TM: tunicamycin), cell-wall stress (CFW: calcofluor white, CR: Congo red, SDS: Sodium dodecyl sulfate, caffeine), osmotic stress (NaCl, KCl, sorbitol) and treatment with antifungal drugs (fluconazole, ketoconazole, fludioxonil). (C) Spotting analysis of *C. neoformans mnl1*Δ, *mnl2*Δ, and *mnl1*Δ*mnl2*Δ cultivated under various stress-inducing conditions.

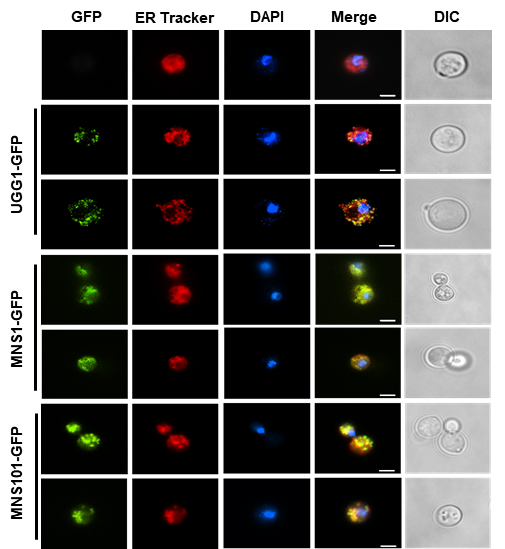

**Supplementary Figure S4.** **Subcellular localization of GFP-tagged Ugg1, Mns1, and Mns101.** ER tracker was used to visualize the ER membrane. Scale bar, 2.5 μm

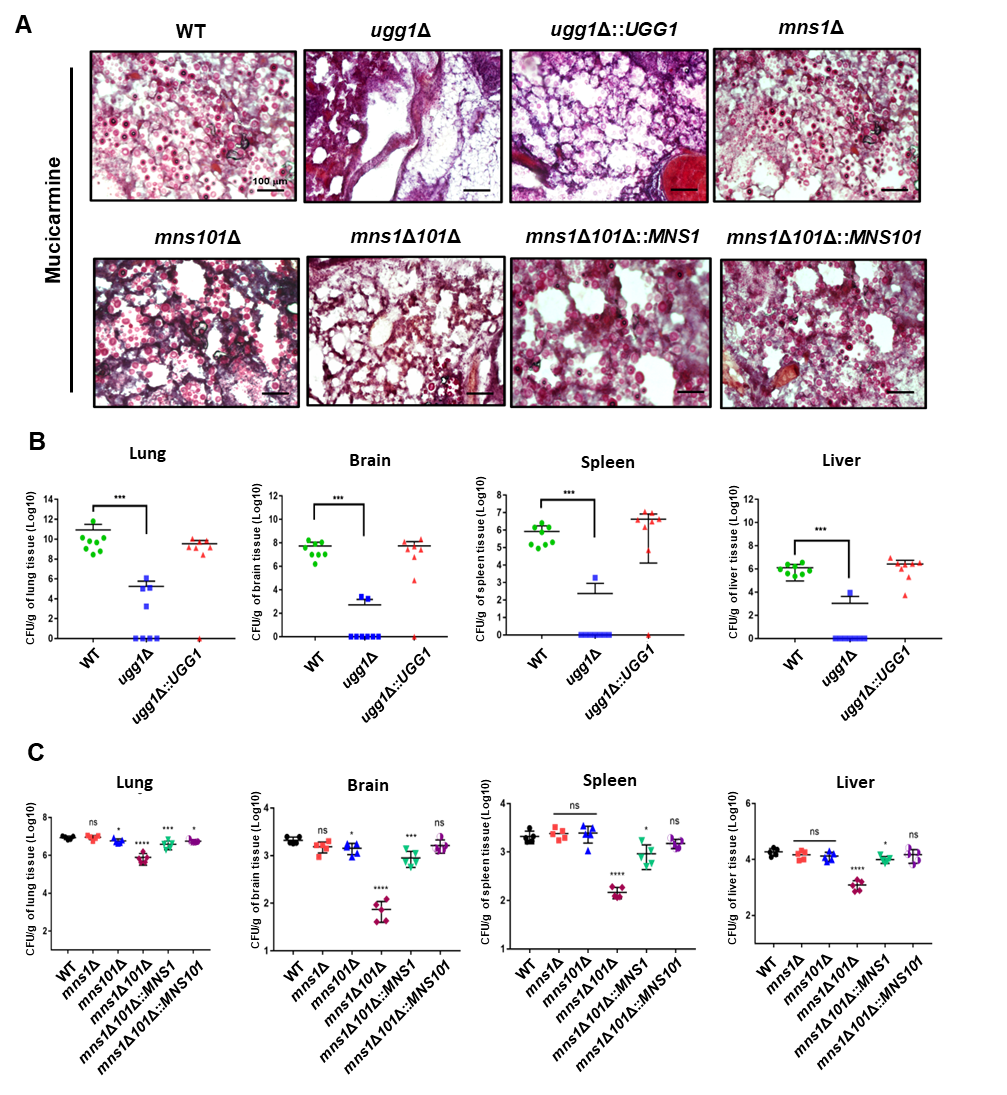

**Supplementary Figure S5.** **Fungal burden of mice infected with *C. neoformans* ERQC-related mutant cells.** (A) Histopathological analysis of lung tissue infected with *C. neoformans*. Lungs were excised at the death point and stained with mucicarmine to visualize *C. neoformans* cells. (B) Distribution of *C. neoformans* cells (wild type (WT), *ugg1*Δ, and *ugg1*Δ::*UGG1*) in systemic organs at the humane endpoint (60 days). The fungal burden was determined using colony forming unit (CFU) counts after plating on YPD medium supplemented with chloramphenicol. (C) Fungal burden assay of WT, *mns1*Δ, *mns1*Δ, and *mns1*Δ*101*Δ at the early stage of infection (7 dpi). Statistical significance: **** *P*<0.0001, *** *P*<0.0005, * *P* < 0.05*,* ns, not significant. All statistical data were determined by one-way ANOVA and Dunnett’s post-hoc test

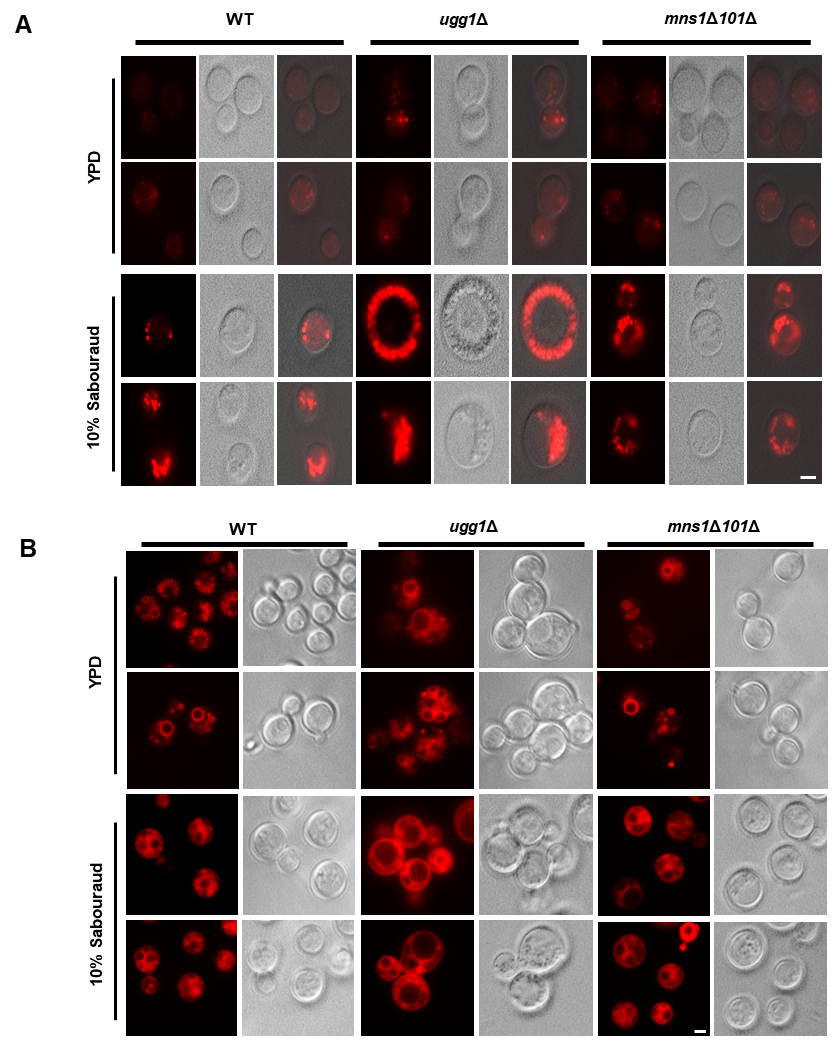

**Supplementary Figure S6.** **Lipid and vacuole staining of WT and *ugg1*Δ strains.** Respective strains were cultivated in either YPD or 10% Sabouraud media at 30 °C and washed twice with PBS. Cells (OD_600_ = 1) were stained with dyes, LipidTox (Invitrogen) for detection of LDs (A) or FM4-64 (Invitrogen) for detection of vacuoles (B), for 30 min at room temperature in the dark. Cells were visualized using an Eclipse Ti-E fluorescence microscope (Nikon) equipped with a Nikon DS-Qi2 camera and a Plan Apo VC 100X oil differential interference contrast (DIC) lens. Images were processed using the NIS-Elements imaging software (Nikon). All images are shown on the same scale. Scale bar, 2.5 µm.

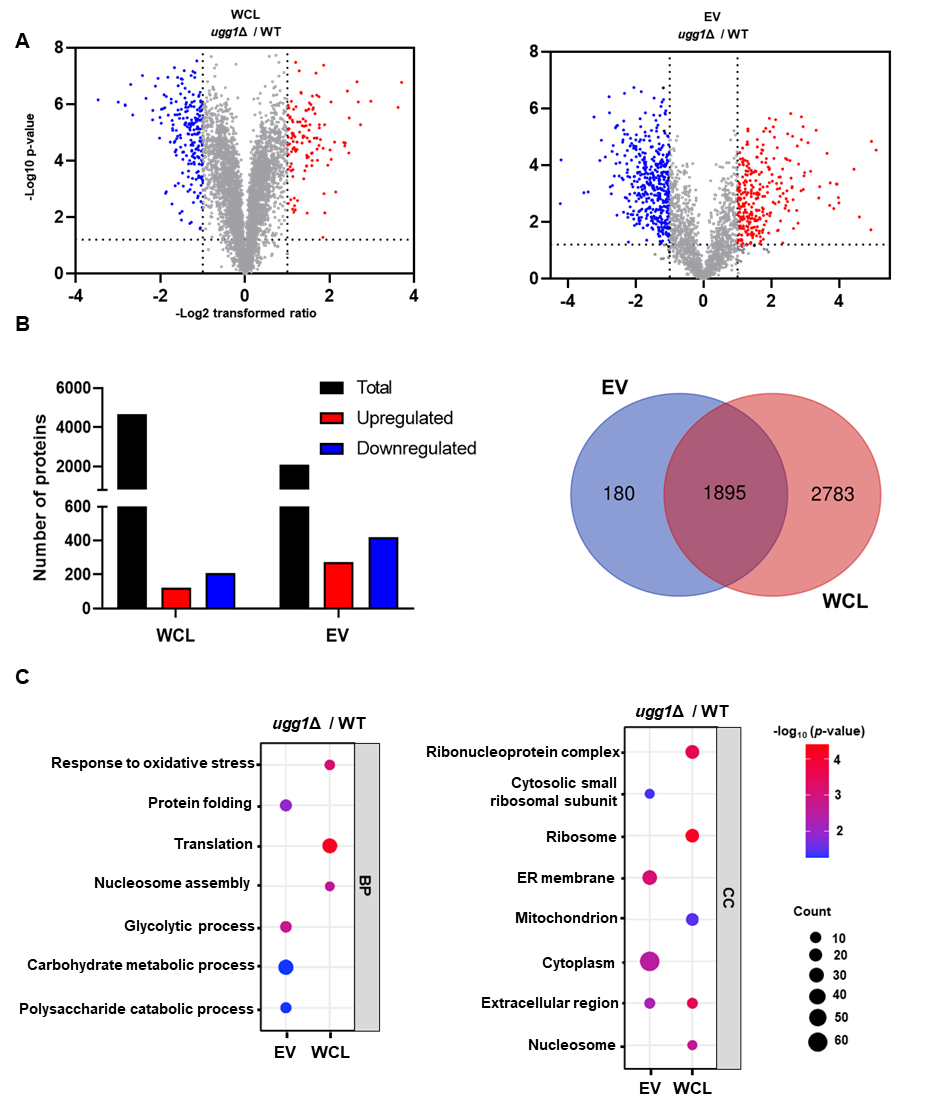
 **Supplementary Figure S7. Comparative proteomic analysis of whole-cell lysate (WCL) and extracellular vesicles (EVs) from WT and *ugg1*Δ strains.** EVs were isolated from *C. neoformans* cells cultivated on synthetic dextrose (SD) solid medium for 24h. (A) Volcano plot highlighting proteins exhibiting a > 2-fold change in expression. (B) Total number of proteins identified in WCL and EV samples, with red and blue columns representing proteins with > 2-fold differential expression. (C) Gene Ontology (GO) enrichment analysis of differentially expressed proteins (> 2-fold), categorized into biological process (BP) and cellular component (CC). Bubble plots were generated using the SRPlot web tool (<http://www.bioinformatics.com.cn/srplot>; Tang et al., 2023).

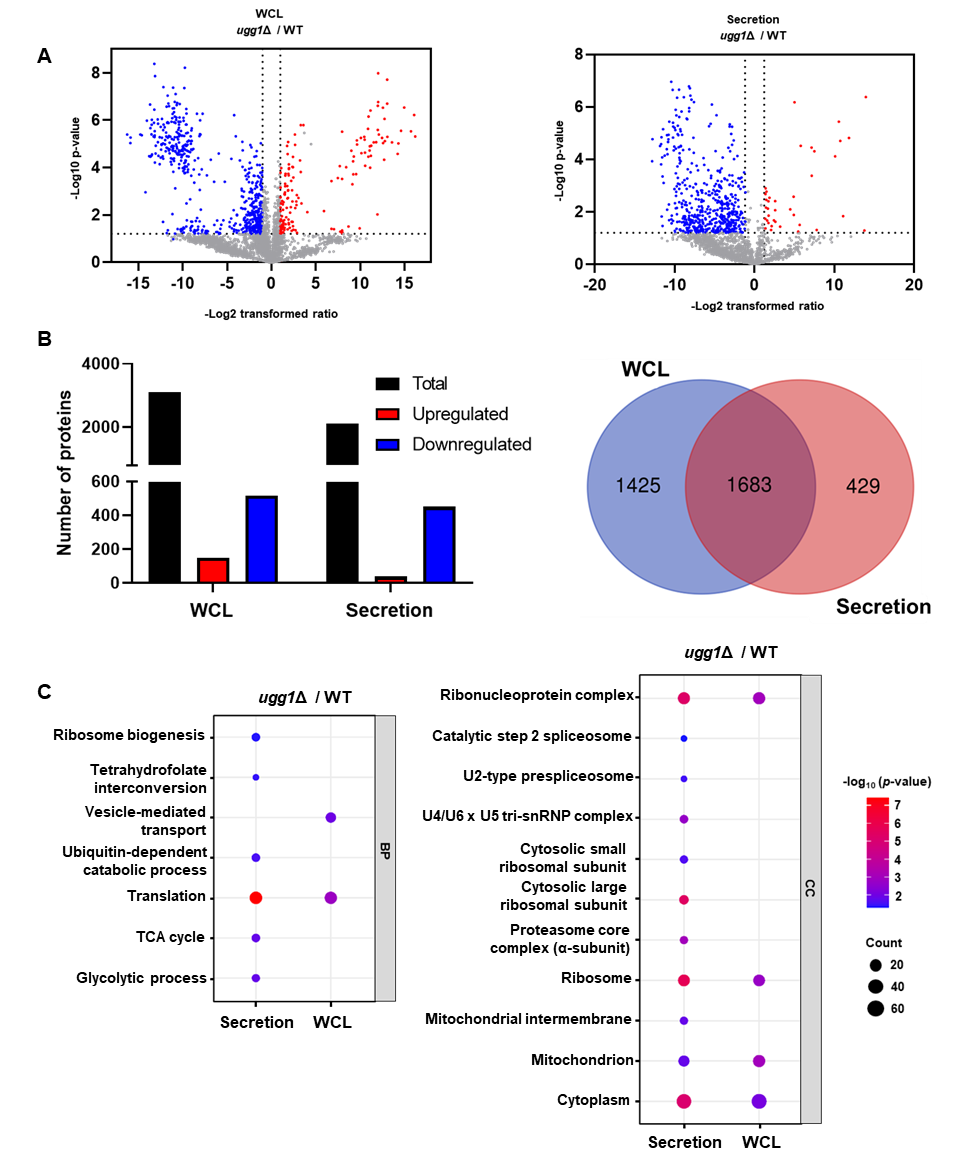

**Supplementary Figure S8. Comparative proteomic analysis of whole-cell lysate (WCL) and secretion fractions from WT and *ugg1*Δ strains.** *C. neoformans* cells were cultivated in the synthetic dextrose (SD) liquid broth for 24 h. Secretome samples were prepared by TCA precipitation of culture supernatants. (A) Volcano plot highlighting proteins exhibiting a > 2-fold change in expression. (B) Total number of proteins identified in WCL and EV samples, with red and blue columns representing proteins with > 2-fold differential expression. (C) Gene Ontology (GO) enrichment analysis of differentially expressed proteins (> 2-fold), categorized into biological process (BP) and cellular component (CC). Bubble plots were generated using the SRPlot web tool (<http://www.bioinformatics.com.cn/srplot>; Tang et al., 2023).
