## Supplementary material for "Evolutionary unique *N*-glycan-dependent protein quality control system plays pivotal roles in cellular fitness and extracellular vesicle transport in *Cryptococcus neoformans*": Table S1

**Supplementary Table S1A. List of strains used in this study**

| **Strain** | **Genotype** | **Parent** | **Reference** |
| --- | --- | --- | --- |
| *C*. *neoformans* H99 | *MATα* (serotype A) |  | Perfect et al., 1993 |
| *ugg1*Δ | *MATα* Cn03648::*NAT*#159 | H99 | This study |
| *ugg1*Δ::*UGGT* | *MATα* Cn03648::*NAT*#159 Cn0348::*NEO* | *ugg1*Δ | This study |
| *mns1*Δ | *MATα* Cn2081::*HyB* | H99 | This study |
| *mns101*Δ | *MATα* Cn3240:: *NAT*#159 | H99 | This study |
| *mns1*Δ::*MNS1* | *MATα* Cn02081::*HyB*Cn02081::*NEO* | *mns1*Δ | This study |
| *mns101*Δ::*MNS101* | *MATα* Cn03240::*NAT*#159Cn03240::*NEO* | *mns101*Δ | This study |
| *mns1*Δ*101*Δ | *MATα* Cn03240::*NAT*#159Cn02081::*HyB* | *mns1*Δ | This study |
| *mns1*Δ*101*Δ::*MNS1* | *MATα*Cn03240::*NAT*#159Cn02081::*HyB*::Cn02081::*NEO* | *mns1*Δ*101*Δ | This study |
| *mns1*Δ*101*Δ::*MNS101* | *MATα*Cn03240::*NAT*#159Cn02081::*HyB*::Cn02081::*NEO* | *mns1*Δ*101*Δ | This study |
| *mnl1*Δ | *MATα* Cn01981::*NAT*#159 | H99 | This study |
| *mnl2*Δ | *MATα* Cn04498::*NAT*#159 | H99 | This study |
| *mnl1*Δ*mnl2*Δ | *MATα* Cn01981::*NAT*#159Cn04498::HyB | *mnl2*Δ | This study |
| *cac1*Δ | *MATα* *cac1*::*NAT*#159 | H99 | Bahn et al., 2004 |
| *cap59*Δ | *MATα* Cn00721::*HYB* | H99 | Thak et al., 2020 |
| *rim101*Δ | *MATα* *rim101*::*NAT* | H99 | O’Meara et al., 2010 |
| *ugg1*Δ::*GFP-UGG1* | *MATα* Cn3648::*NAT*#159SP_CnGFP_Cn03648::*NEO* | *ugg1*Δ | This study |
| *MNS1-GFP* | *MATα* Cn02081-CnGFP::*NEO* | H99 | This study |
| *MNS101-GFP* | *MATα* Cn03240-CnGFP::*NEO* | H99 | This study |

*Each *NAT-STM#* indicates the Nat^r^ marker with a unique signature tag

**Supplementary Table S1B. List of plasmids used in this study**

| **Plasmid** | **Description** | **Reference** |
| --- | --- | --- |
| pNAT-STM#159 | NAT-resistant marker vector for gene disruption | Kim et al., 2009 |
| pJAF | pJAF-based vector containing hygromycin B marker | Hua et al., 2000 |
| pJAFS1 | NEO-resistant marker vector | Cheon et al., 2011 |
| pWH091 | NEO-resistant marker vector containing codon optimized GFP for *C. neoformans* (Cn*GFP*) | Jung et al., 2009 |
| pT-CnUGG1D_L | pT-Blunt^TM^-based vector containing the CnUGG1D_L fragment | This study |
| pT-CnUGG1D_R | pT-Blunt^TM^-based vector containing the CnUGG1D_R fragment | This study |
| pJAFS1_UGG1Com | pJAFS1 containing the *UGG1* ORF | This study |
| pT-CnMNS1D_L | pT-Blunt^TM^-based vector containing the CnMNS1D_L fragment | This study |
| pT-CnMNS1D_R | pT-Blunt^TM^-based vector containing the CnMNS1D_R fragment | This study |
| pT-CnMNS101D_L | pT-Blunt^TM^-based vector containing the CnMNS101D_L fragment | This study |
| pT-CnMNS101D_R | pT-Blunt^TM^-based vector containing the CnMNS101D_R fragment | This study |
| pJAFS1_MNS1Com | pJAFS1 containing the *MNS1* ORF | This study |
| pJAFS1_MNS101Com | pJAFS1 containing the *MNS101* ORF | This study |
| pT-CnMNL1D_L | pT-Blunt^TM^-based vector containing the CnMNL1D_L fragment | This study |
| pT-CnMNL1D_R | pT-Blunt^TM^-based vector containing the CnMNL1D_R fragment | This study |
| pT-CnMNL2D_L | pT-Blunt^TM^-based vector containing the CnMNL2D_L fragment | This study |
| pT-CnMNL2D_R | pT-Blunt^TM^-based vector containing the CnMNL2D_R fragment | This study |
| pJAFS1_SP_CnGFP_UGG1 | pJAFS1 containing the Cn*GFP-UGG1* fusion | This study |
| pWH091-MNS1 | pWH091 containing the *MNS1*-Cn*GFP* fusion | This study |
| pWH091-MNS101 | pWH091 containing the *MNS101-*Cn*GFP* fusion | This study |

**Supplementary Table S1C. List of primers used in this study.**

| **Name** | **Sequence (5′-3′)** | **Purpose** |
| --- | --- | --- |
| 03648p_Fw | TTGCAGCGCTTATTTCCC | *UGG1* disruption cassette |
| 03648p_Rv | GCTCACTGGCCGTCGTTTTACCCCGTTCTATTGTGCAGG | *UGG1* disruption cassette |
| 03648t_Fw | CATGGTCATAGCTGTTTCCTGGTCGACCTTTGTCAAAACCC | *UGG1* disruption cassette |
| 03648t_Rv | GCTGAACGCTACTCACTTGA | *UGG1* disruption cassette |
| M13Fe | GTAAAACGACGGCCAGTGAGC | Screening for selection marker genes (*NAT/NEO*) |
| NSL-2 | AACTCCGTCGCGAGCCCCATCAAC | 5′-Region of *NAT* split marker |
| M13Re | CAGGAAACAGCTATGACCATG | Screening for selection marker genes (*NAT/NEO*) |
| NSR-2 | AAGGTGTTCCCCGACGACGAATCG | 3′-Region of *NAT* split marker |
| UGG1 deletion check L | ATGAGAGCAAGCACAGTAGC | Screening for the *ugg1*Δ strain |
| UGG1 deletion check R | GTTACTCTCATTCCGAGCC | Screening for the *ugg1*Δ strain |
| 03648 NAT integration L | TTTACGACAGCGTGGCCCTA | Screening for the *ugg1*Δ strain |
| 03648 NAT integration R | CTAGCACCCATGATCCAATG | Screening for the *ugg1*Δ strain |
| UGG1 promoter Fw | GGGGGCGGCCGCTAGAACGTGCGACCACCGTC | *UGG1* complementation cassette |
| UGG1 promoter Rv | GGGGGATATCCGACGACGGACTTTGAGAAG | *UGG1* complementation cassette |
| UGG1 infusion Left Fw | GTCGACCTCGAGGGGGGGCCC*GATATC*GCTCCAAGACGAGGAAGATG | *UGG1* complementation cassette |
| UGG1infusion Left Rv | CCGCCAAACGTCATACTGTG | *UGG1* complementation cassette |
| UGG1infusion right Fw | CACAGTATGACGTTTGGCGG | *UGG1* complementation cassette |
| 02081_L1 | CTGTGGAGATCTCCTCGAT | *MNS1* disruption |
| 02081_L2 | GCTCACTGGCCGTCGTTTTACGCGGTCTTGCGATGTGTT | *MNS1* disruption |
| 02081_R1 | CATGGTCATAGCTGTTTCCTGCTCCCAGCAACGGTATCCT | *MNS1* disruption cassette |
| 02081_R2 | GTCTCAAAGCTGATGTCTGC | *MNS1* disruption cassette |
| 02081_del check L | GGGCAGTTTGTGGCTTCTG | Screening for the *mns1*Δ strain |
| 02081_del check R | GACCCTACACCACTTCTGG | Screening for the *mns1*Δ strain |
| 02081_integration_F1 | AAAGTTCGACAGCGTCTC | Screening for the *mns1*Δ strain |
| 02081_integration_R1 | CCCAAGCTGCATCATCGAAA | Screening for the *mns1*Δ strain |
| mns1 comp F1 | ATCGATACCGTCGACCTCGAGCAATAGCCGACGGTAGTC | *MNS1* complementation cassette |
| mns1 comp R1 | TGGGAGCAATACCATCATGG | *MNS1* complementation cassette |
| mns1 comp F2 | CCATGATGGTATTGCTCCCA | *MNS1* complementation cassette |
| mns1 comp R2 | GGTACCGGGCCCCCCCTCGAGTGGTTAACCAGGCTGCTC | *MNS1* complementation cassette |
| ORFconfirm_Fr | CAATAGCCGACGGTAGTC | Screening for *MNS1* complementation |
| ORFconfirm_Rv | TCAGTTATGTCACCCCCG | Screening for *MNS1* complementation |
| NEOinteg F1 | TGGATTGCACGCAGGTTCT | Screening for *MNS1* complementation |
| NEOinteg R1 | GAAGAACTCGTCAAGAAGGC | Screening for *MNS1* complementation |
| 03240_L1 | CGGATCACCACCAGATCACC | *MNS101* disruption cassette |
| 03240_L2 | GCTCACTGGCCGTCGTTTTACGGTATGTGGGGGGCTTGACA | *MNS101* disruption cassette |
| 03240_R1 | CATGGTCATAGCTGTTTCCTGATCCCATCGCTGGTCGATC | *MNS101* disruption cassette |
| 03240_R2 | CGCCGTTTGGAACCACGAT | *MNS101* disruption cassette |
| 03240_deletion_L | ATGGGTCGCTCTCTTTCC | Screening for the *mns101*Δ strain |
| 03240_deletion_R | TCGGCAAGACCACCAAAG | Screening for the *mns101*Δ strain |
| 03240_NAT integration_L | CCCCCTCTCTAAGTACAA | Screening for the *mns101*Δ strain |
| 03240_NAT integration_R | ACTCAGTCCTCCATCCTCCA | Screening for the *mns101*Δ strain |
| mns101_comp_F1 | ATCGATACCGTCGACCTCGAGGGTATTCTCTCGTTCCGC | *MNS101*complementation cassette |
| mns101_comp_R1 | TGTACCAGGGTCCATTCG | *MNS101*complementation cassette |
| mns101_comp_F2 | CGAATGGACCCTGGTACA | *MNS101*complementation cassette |
| mns101_comp_R2 | GGTACCGGGCCCCCCCTCGAGCATGTCTCCTCCTCATTC | *MNS101*complementation cassette |
| MNS101comp_confirmFr | CATGAGGTACGACCACCT | Screening for *MNS101* complementation |
| MNS101comp_confirmRv | CCCAGGATCCATTTAGGC | Screening for *MNS101* complementation |
| 01987_L1 | AGGTTGCACAGATGCATAGC | *MNL1* disruption cassette |
| 01987_L2 | CGATTCGCGGACTAGGGAA | *MNL1* disruption cassette |
| 01987_R1 | TGGTCCTTGATAATAGCAT | *MNL1* disruption cassette |
| 01987_R2 | ACAAATGGACAAGAGGCGGT | *MNL1* disruption cassette |
| 01987_deletion_L | CTCTCTAGATTGACAGGCG | Screening for the *mnl1*Δ strain |
| 01987_deletion_R | TACGCGAGAGCACTCGTCTT | Screening for the *mnl1*Δ strain |
| 01987_NAT integration_L | CGGCCAAGTAACTGGTTTTC | Screening for the *mnl1*Δ strain |
| 01987_NAT integration_R | ATCGTTGTTGGGCTTGGG | Screening for the *mnl1*Δ strain |
| 04498_L1 | CTGGAGCACTCAAACTGAC | *MNL2* disruption cassette |
| 04498_L2 | GCTCACTGGCCGTCGTTTTACGAGACTCGCAATTCGCAA | *MNL2* disruption cassette |
| 04498_R1 | CATGGTCATAGCTGTTTCCTGACAGAGGGCGATCATTGG | *MNL2* disruption cassette |
| 04498_R2 | TGACGGAAGCAGGTGAGTCT | *MNL2* disruption cassette |
| 04498_deletion_L | GTGCCCATACAACCAAACG | Screening for the *mnl2*Δ strain |
| 04498_deletion_R | GACAAACATGTGCTGGGC | Screening for the *mnl2*Δ strain |
| 04498_HyB integration_L | TGACCTTGCGCGCATGAA | Screening for the *mnl2*Δ strain |
| 04498_ HyB integration_R | TGGAGGATGGAGGACTGA | Screening for the *mnl2*Δ strain |
| C17 | ATGGCTACCGCTGTCGCT | RT-PCR primer for *Hxl1*; Cheon et al., 2011 |
| C18 | TGATTCGCGGTTACGGAT | RT-PCR primer for *Hxl1*; Cheon et al., 2011 |
| C19 | CACTCCATTCCTTTCTGC | RT-PCR primer for *Hxl1*; Cheon et al., 2011 |
| C20 | CGTAACTCCACTGTGTCC | RT-PCR primer for *Hxl1*; Cheon et al., 2011 |
| C27 | TCGATGCCAATGGTATCC | RT-PCR primer for *KAR2*; Cheon et al., 2011 |
| C28 | TCATGGCTGAAAGGCATC | RT-PCR primer for *KAR2*; Cheon et al., 2011 |
| C33 | AGCCTTCTCTCCTTGGTC | RT-PCR primer for *ACT1*; Cheon et al., 2011 |
| C34 | ACGATTGAGGGACCAGAC | RT-PCR primer for *ACT1*; Cheon et al., 2011 |
| C51 | CTTCCAGCCTTCTCTCCTTG | qRT-PCR primer for *ACT1*; Cheon et al., 2011 |
| C52 | AGAGGTCCTTCCTGATGTCG | qRT-PCR primer for *ACT1*; Cheon et al., 2011 |
| C53 | CTCTGAGGACGACAAGGACA | qRT-PCR primer for *KAR2*; Cheon et al., 2011 |
| C54 | AGCTCAGAAAGCTGCTCCTC | qRT-PCR primer for *KAR2*; Cheon et al., 2011 |
| Bv_Not1_ugg1_pro _F1 | AGCTCCACCGCGGTGGCGGCCGCTAGAACGTGCGACCA | GFP-Ugg1 fusion construct |
| SP_03648pro_R1 | CTGAAGCTGCGAGGGCTAGAGCTACTGTGCTTGCTCTCATAATGCCCGTTCTATT | GFP-Ugg1 fusion construct |
| SP_CnGFP_F2 | TCTAGCCCTCGCAGCTTCAGCCCTAGCGGCGTCCGCGTCTGTGAGCAAGGGCGAG | GFP-Ugg1 fusion construct |
| UGG1ORF_5gly_CnGFP_R2 | AGACTTACGCGTACTGGTGGACCGCCACCGCCACCGGACTTGTACAGCTCGTCCA | GFP-Ugg1 fusion  construct |
| UGG1ORF_5gly_CnGFP_F3 | TGGACGAGCTGTACAAGTCC*GGTGGCGGTGGCGGT*CCACCAGTACGCGTAAGTCT | GFP-Ugg1 fusion construct |
| Bv_Mfe1_UGG1ORF _R3 | GAGGATGGATAGGCCAATTGTGATACCTTTTGTCTTTCTC | GFP-Ugg1 fusion construct |
| BamHI_MNS1 F1 | CTTGGTACCGAGCTCGGATCCTAGAGGCGCCATCGAAGCAAAG | Mns1-GFP fusion construct |
| MNS1_GFP R1 | CTCGCCCTTGCTCACACCGCCACCGCCACCGGATCCTGAAAGGGCGAAAGAAGAG | Mns1-GFP fusion construct |
| BamHI_ MNS101 F1 | CTTGGTACCGAGCTCGGATCCTGATGGAAGAGATCAACATGTT | Mns101-GFP fusion construct |
| MNS101_GFP R1 | CTCGCCCTTGCTCACACCGCCACCGCCACCGGATCCATCGACCTTATTTTTTAAAACCTCTGG | Mns101-GFP fusion construct |
| qRT_UGG1_Fw | CGACCTTCCCTCCAACCC | qRT-PCR primer for *UGG1* |
| qRT_UGG1_Rv | GGAAGGTGACAGCGACAGG | qRT-PCR primer for *UGG1* |
| qRT_MNS1_Fw | ATCTATGCCGCCCAAGCAGT | qRT-PCR primer for *MNS1* |
| qRT_MNS1_Rv | CTGACCGGCCTCGAGCTAAA | qRT-PCR primer for *MNS1* |
| qRT_ MNS101_Fw | CATCGGCGCTATGCTTGGTC | qRT-PCR primer for *MNS101* |
| qRT_ MNS101_Rv | ACATGGCCGAGGTCTCATGG | qRT-PCR primer for *MNS101* |
| qRT_MNL1_Fw | CCTACGTCCAACGAGACGCT | qRT-PCR primer for *MNL1* |
| qRT_MNL1_Rv | TTGCCCGTCCAGTCTCCATC | qRT-PCR primer for *MNL1* |
| qRT_MNL2_Fw | ATCGCCTCCCATCTCTCCCT | qRT-PCR primer for *MNL2* |
| qRT_MNL2_Rv | TGATCGCCCTCTGTCCAACC | qRT-PCR primer for *MNL2* |
| qRT_GAPDH_Fw | CCGCTAACATCATCCCTTCT | qRT-PCR primer for *GAPDH* |
| qRT_GAPDH_Rv | CCACGACGGATACATCAGAG | qRT-PCR primer for *GAPDH* |
| qRT_Cap60_Fw | GCTCACGAGGGTGGAAACT | qRT-PCR primer for *CAP60* |
| qRT_Cap60_Rv | GCCTACTCTTCTCTGGCTC | qRT-PCR primer for *CAP60* |
| qRT_Cap59_Fw | TTGGGACGGTGCTGGTGAT | qRT-PCR primer for *CAP59* |
| qRT_Cap59_Rv | CCATCCAGGCATATTTCGG | qRT-PCR primer for *CAP59* |
| qRT_Cap64_Fw | CCGTCCCAGTGATATCCTCA | qRT-PCR primer for *CAP64* |
| qRT_Cap64_Rv | CGGCTCCTTTACTTGGGTG | qRT-PCR primer for *CAP64* |
| qRT_Cap2_Fw | CACCGGATGAGACTGCATCT | qRT-PCR primer for *CAP2* |
| qRT_Cap2_Rv | CCCTCTCCCATCTTCTTACG | qRT-PCR primer for *CAP2* |
| qRT_Cap10_Fw | TGCTCGATGCCTCCAATTCC | qRT-PCR primer for *CAP10* |
| qRT_Cap10_Rv | TGAACATCCACTCCCTACCC | qRT-PCR primer for *CAP10* |
| qRT_Pmt4_Fw | GACCACAAGACTACGGCCA | qRT-PCR primer for *PMT4* |
| qRT_Pmt4_Rv | GGCCATACGTCAATGGACTC | qRT-PCR primer for *PMT4* |
| qRT_Cas1_Fw | AGGTCGAACAGCAGAAGGGT | qRT-PCR primer for *CAS1* |
| qRT_Cas1_Rv | ACCGCAATCTTGACCGAGGT | qRT-PCR primer for *CAS1* |
| qRT_Uge1_Fw | GGCTGCTACCATTCCACTCA | qRT-PCR primer for *UGE1* |
| qRT_Uge1_Rv | GGGAAGTACTCCTTTGCCTC | qRT-PCR primer for *UGE1* |
| qRT_Ugd1_Fw | GCCCAGCGAATTTCCTCTG | qRT-PCR primer for *UGD1* |
| qRT_Ugd1_Rv | CGCCTTGAGGAACTTGGAAC | qRT-PCR primer for *UGD1* |
| qRT_Chs4_Fw | GGGTGACCAGAGTCGTATCA | qRT-PCR primer for *CSH4* |
| qRT_Chs4_Rv | TCCAGCACCTCCTCATCTTG | qRT-PCR primer for *CSH4* |
| qRT_Chs7_Fw | TCCATACTTGAGTGGGGGTC | qRT-PCR primer for *CSH7* |
| qRT_Chs7_Rv | CTTTTTGATGGTAGCGCCGC | qRT-PCR primer for *CSH7* |
| qRT_Skn1_Fw | GGATCAGCCGGGAGATAGTA | qRT-PCR primer for *SKN1* |
| qRT_Skn1_Rv | ATACCCCTGATCCCTTGCCT | qRT-PCR primer for *SKN1* |
| qRT_Kre6_Fw | GGCGACTCATTCCCCAAGAA | qRT-PCR primer for *KRE6* |
| qRT_Kre6_Rv | TCCGATCGTTGGTACACACG | qRT-PCR primer for *KRE6* |
| qRT_CNAG_06336_Fw | GTCGCTGCAGAAGCCTCTAA | qRT-PCR primer for CNAG_06336 |
| qRT_CNAG_06336_Rv | GCTAGCATCACCTTGAGGG | qRT-PCR primer for CNAG_06336 |
| qRT_Exg104_Fw | CGGCCTAGCTCTTGTCATTC | qRT-PCR primer for *EXG104* |
| qRT_Exg104_Rv | TCCAAGCGACATACTCGTCG | qRT-PCR primer for *EXG104* |
| qRT_Ebg1_Fw | GCCGACGCTGTTTGGAACTA | qRT-PCR primer for *EBG1* |
| qRT_Ebg1_Rv | GGCTTCAGTAAAGGCAGAGC | qRT-PCR primer for *EBG1* |
| qRT_Sav1_Fw | TGAACAACATGCATCGCCCG | qRT-PCR primer for *SAV1* |
| qRT_Sav1_Rv | AACCGCAGTCCAAACTCGTC | qRT-PCR primer for *SAV1* |
| qRT_Sec6_Fw | GCCCGTCGCCATTCAAAAGA | qRT-PCR primer for *SEC6* |
| qRT_Sec6_Rv | GAGATGCCGCGAGCATGTT | qRT-PCR primer for *SEC6* |
| qRT_Sec14_Fw | CTCGACATCCCCAAGCTCTA | qRT-PCR primer for *SEC14* |
| qRT_Sec14_Rv | GTGTCCAACCTCCTCGGAA | qRT-PCR primer for *SEC14* |
| qRT_Arf1_Fw | CAGGTATCCGAGGCTCTCAA | qRT-PCR primer for *ARF1* |
| qRT_Arf1_Rv | CGCGCGTCTAACTTTGTGAC | qRT-PCR primer for *ARF1* |
| qRT_Vph1_Fw | GTTTGACTTTAACGGCGGGC | qRT-PCR primer for *VPH1* |
| qRT_ Vph1_Rv | CCCAATGCAACCTCAATGCG | qRT-PCR primer for *VPH1* |
| qRT_Apt1_Fw | CAAAACCATGGCCAGCACGA | qRT-PCR primer for *APT1* |
| qRT_ Apt1_Rv | CCCCGAAACCTCTCCCTAAA | qRT-PCR primer for *APT1* |
| qRT_Grasp_Fw | CGCTTGGTCAGAAGAAGCGT | qRT-PCR primer for *GRASP* |
| qRT_Grasp_Rv | GGCTCCCTTCTAAGACATCC | qRT-PCR primer for *GRASP* |
| qRT_Aim25_Fw | GGCATGACCGAGGATGAAC | qRT-PCR primer for *AIM25* |
| qRT_Aim25_Fw | ACCCTTCCAAACCTTCTGGG | qRT-PCR primer for *AIM25* |
| qRT_Cin1_Fw | TAACGGCTCCAACGGCGTT | qRT-PCR primer for *CIN1* |
| qRT_Cin1_Rv | CCTCGAGAAGGACGATGA | qRT-PCR primer for *CIN1* |
| qRT_Vps15_Fw | TCGAATGCCCGAAGGTTCAC | qRT-PCR primer for *VPS15* |
| qRT_Vps15_Rv | ATCGCGTCATAGTGGGGTTG | qRT-PCR primer for *VPS15* |
| qRT_Vps27_Fw | TCAGGAGGTGTCAGTGGCT | qRT-PCR primer for *VPS27* |
| qRT_Vps27_Rv | CCCTGATACCTTCCTTTCCC | qRT-PCR primer for *VPS27* |
| qRT_Vps34_Fw | TCCAGAGCCTATGCTACACC | qRT-PCR primer for *VPS34* |
| qRT_Vps34_Rv | GCTTTATCTGGCTCCAGCTG | qRT-PCR primer for *VPS34* |
| qRT_Hse1_Fw | CGTATTATCCAGCCCATGCG | qRT-PCR primer for *HSE1* |
| qRT_Hse1_Rv | CCGACTGTCACACCTTCCA | qRT-PCR primer for *HSE1* |
| qRT_CNAG_07029_Fw | CCTCCTCGTGGACAAGACA | qRT-PCR primer for CNAG_07029 |
| qRT_CNAG_07029_Rv | AGGCCAACAACACCGGATAG | qRT-PCR primer for CNAG_07029 |

^*^Underlined sequences correspond to restriction enzyme sites
