## Supplementary material for "Evolutionary unique *N*-glycan-dependent protein quality control system plays pivotal roles in cellular fitness and extracellular vesicle transport in *Cryptococcus neoformans*": Table S2

**Supplementary Table S2A. Upregulated (> 2-fold) *Cryptococcus neoformans* genes in** ***ugg1*Δ over wild type (WT) cultivated in yeast extract peptone dextrose (YPD) at 30 °C**

| **Locus tag (CNAG)** | **Gene name** | ***uggt*Δ/WT**  **(Fold change)** | **Description** |
| --- | --- | --- | --- |
| **Hydrolyzing O-glycosyl compounds in cell wall** | | | |
| CNAG_00897 | *SKN1* | 3.133 | Glucosidase for beta-1,6-glucan synthesis |
| CNAG_00914 | *KRE6* | 2.164 | Glucosidase for beta-1,6-glucan synthesis |
| CNAG_02860 | *EBG1* | 2.248 | Endo-1,3(4)-beta-glucanase |
| CNAG_04874 |  | 14.898 | Hypothetical protein |
| CNAG_05411 | *LPI9* | 2.097 | Endoglucanase |
| CNAG_05458 |  | 10.821 | Endo-1,3(4)-beta-glucanase |
| CNAG_06336 | *BLG2* | 5.195 | Glucan 1,3-beta-glucosidase |
| **Carbohydrate metabolic process** | | | |
| CNAG_00546 | *CHS4* | 2.024 | Chitin synthase |
| CNAG_00744 | *OCH1* | 2.202 | Alpha-1,6-mannosyltransferase |
| CNAG_00799 |  | 2.214 | Cellulase |
| CNAG_01230 | *CDA2* | 2.473 | Chitin deacetylase 2 |
| CNAG_01954 |  | 4.953 | Aldo-keto reductase |
| CNAG_02189 | *Amy1* | 2.406 | Alpha-amylase |
| CNAG_02217 | *CHS7* | 27.261 | Chitin synthase export chaperone |
| CNAG_04944 |  | 2.489 | Hypothetical protein |
| CNAG_05264 | *AmyA* | 6.331 | Alpha-amylase |
| CNAG_05828 | *QRI1* | 2.19 | UDP-N-acetylglucosamine pyrophosphorylase |
| CNAG_06033 | *MAK3202* | 2.223 | pfkB family carbohydrate kinase superfamily |
| CNAG_06898 | *CHS7* | 2.326 | Chitin synthase export chaperone |
| **Catalytic activity in membrane traffic** | | | |
| CNAG_04655 |  | 2.432 | Rab family GTP-binding protein |
| **DNA replication, cell cycle and differentiation** | | | |
| CNAG_00498 | *CDC14* | 2.175 | Cell division cycle protein 14 |
| CNAG 03366 | *ZNF2* | 2.271 | Transcription factor regulating hyphal growth |
| CNAG_04696 |  | 2.436 | DNA clamp loader |
| CNAG_07406 |  | 5.6 | Pheromone alpha |
| CNAG_07970 | *EME1* | 2.469 | Crossover junction endonuclease |
| **Integral component of membrane** | | | |
| CNAG_00668 |  | 2.016 | Hypothetical protein |
| CNAG_01668 |  | 8.304 | Hypothetical protein |
| CNAG_01713 |  | 2.543 | Hypothetical protein |
| CNAG_02156 |  | 2.98 | Hypothetical protein |
| CNAG_02220 |  | 5.111 | Hypothetical protein |
| CNAG_02342 |  | 2.116 | Hypothetical protein |
| CNAG_02921 |  | 2.105 | Hypothetical protein |
| CNAG_03647 |  | 2.252 | Hypothetical protein |
| CNAG_03857 |  | 4.364 | Hypothetical protein |
| CNAG_05654 | *RIM90* | 2.066 | Hypothetical protein, SUR7/PalI family |
| CNAG_06000 | *CMP1* | 2.568 | Putative mannoprotein |
| CNAG_07943 |  | 6.601 | Hypothetical protein |
| CNAG_07981 |  | 4.385 | Hypothetical protein |
| **Lipid metabolic process** | | | |
| CNAG_02008 | *OVA1* | 2.938 | Lipid binding protein |
| CNAG_07638 |  | 2.146 | Hypothetical protein |
| **Oxidoreductase activity** | | | |
| CNAG_00876 | *FRE7* | 3.193 | Ferric-chelate reductase |
| CNAG_01542 | *TauD* | 2.046 | Taurine catabolism dioxygenase |
| CNAG_02049 | *PUT1* | 2.637 | Proline dehydrogenase |
| CNAG_02182 | *GRE2* | 2.336 | D-lactaldehyde dehydrogenase |
| CNAG_02602 |  | 2.716 | Flavonol synthase |
| CNAG_03465 | *LAC1* | 2.073 | Laccase |
| CNAG_04307 | *URO1* | 2.126 | Urate oxidase |
| CNAG_04313 |  | 2.221 | NADPH dehydrogenase |
| CNAG_05256 | *CAT2* | 4.651 | Catalase |
| CNAG_05258 | *SMG1* | 23.15 | Glucose-methanol-choline oxidoreductase |
| CNAG_07491 |  | 2.842 | Glutaredoxin |
| **Proteolysis** | | | |
| CNAG_00919 | *CXD1* | 2.542 | Carboxypeptidase D |
| CNAG_04380 |  | 2.91 | Aspartic peptidase |
| CNAG_04635 |  | 4.596 | Endopeptidase |
| CNAG_06365 | *DER1* | 2.048 | Derlin-2/3 in ER-associated degradation |
| CNAG_06658 |  | 3.66 | Rhomboid family membrane protein for intramembrane proteolysis |
| CNAG_06693 |  | 2.392 | Hypothetical protein |
| **Transporter activity** | | | |
| CNAG_00895 | *ZIP1* | 2.099 | Solute carrier family 39 (zinc transporter), member 1/2/3 |
| CNAG_01174 |  | 2.821 | Hypothetical protein |
| CNAG_04474 |  | 2.322 | Monocarboxylic acid transporter |
| CNAG_05075 |  | 15.295 | Solute carrier family 20 |
| CNAG_05592 | *ECA1* | 2.467 | ER Ca^2+^-ATPase, fungal-type Ca^2+^pump |
| CNAG_06034 |  | 2.227 | Allantoin permease |
| **Ribosome biogenesis and RNA processing** | | |  |
| CNAG_00706 | *RNP24* | 2.073 | RNA-binding protein |
| CNAG_01098 |  | 2.35 |  |
| CNAG_02185 |  | 2.125 | Nuclear protein |
| CNAG_03302 | *LTV1* | 2.626 | Ribosome biogenesis factor |
| CNAG_03354 |  | 3.777 | Hypothetical protein |
| CNAG_03645 |  | 4.273 | NET1-associated nuclear protein 1 (U3 small nucleolar RNA-associated protein 17) |
| CNAG_05228 |  | 2.807 | U3 small nucleolar ribonucleoprotein Lcp5 |
| CNAG_05473 |  | 2.091 | Hypothetical protein |
| CNAG 05755 | *MAK16* | 2.597 | 25S and 5.8S rRNA maturation |
| CNAG_06007 |  | 3.052 | U3 small nucleolar RNA-associated protein 23 |
| CNAG_06340 |  | 2.596 | Pre-rRNA-processing protein TSR4 |
| CNAG_06748 |  | 2.214 | U3 small nucleolar RNA-associated protein 7 |
| CNAG_07413 | *NOG2* | 2.251 | Nucleolar GTP-binding protein 2 |
| **Protein folding** |  |  |  |
| CNAG_02440 | *GRP94* | 2.188 | ER glucose-regulated protein |
| CNAG_03176 | *ERO1* | 3.452 | Endoplasmic oxidoreductin 1 |
| CNAG_03899 | *LSH1* | 2.025 | Chaperone of heat shock protein 70 |
| CNAG_05252 | SCJ1 | 2.31 | Chaperone regulator |
| CNAG_06240 | PDI1 | 2.134 | Protein disulfide-isomerase |
| CNAG_06443 | *KAR2* | 2.309 | Glucose-regulated protein |
| **Serine/threonine kinase activity** | | | |
| CNAG_01704 | *IRK6* | 2.643 | Serine/threonine protein kinase |
| CNAG_02658 | *PCL5* | 2.419 | Regulation of cyclin-dependent serine/threonine kinase |
| CNAG_03385 | *PCL1* | 2.254 | G1/S phase-specific cyclin |

*Genes are grouped based on their Gene Ontology (GO) term or predicted biological functions

**Supplementary Table S2B. Downregulated (> 2-fold) *Cryptococcus neoformans* genes in** ***ugg1*Δ over wild type (WT) cultivated in yeast extract peptone dextrose (YPD) at 30 °C**

| **Locus tag (CNAG)** | **Gene name** | ***uggt*Δ/WT (Fold change)** | **Description** |
| --- | --- | --- | --- |
| **Hydrolyzing O-glycosyl compounds in cell wall** | | | |
| CNAG_00393 | *GLC3* | 0.396 | 1,4-alpha-glucan-branching enzyme |
| CNAG_05913 |  | 0.365 | Alpha-glucosidase |
| **Carbohydrate metabolic process** | | | |
| CNAG_00162 | *AOX1* | 0.247 | Alternative oxidase, mitochondrial |
| CNAG_00866 | *TKL1* | 0.392 | Transketolase |
| CNAG_00984 |  | 0.37 | Glucose and ribitol dehydrogenase |
| CNAG_01078 | *ALD5* | 0.493 | Aldehyde dehydrogenase (NAD) |
| CNAG_02814 |  | 0.486 | Glycerol-3-phosphate dehydrogenase |
| CNAG_02986 | *YSA1* | 0.357 | ADP-ribose pyrophosphatase |
| CNAG_03648 | *KRE5 (UGG1)* | 0.224 | UDP-glucose:glycoprotein glucosyltransferase |
| CNAG_04108 | *PKP2* | 0.485 | Pyruvate dehydrogenase kinase |
| CNAG_04467 |  | 0.492 | Succinate-semialdehyde dehydrogenase (NADP) |
| CNAG_04621 | *GSY1* | 0.5 | Glycogen(starch) synthase |
| CNAG_04659 | *PDC1* | 0.345 | Pyruvate decarboxylase |
| CNAG_04744 | *PMI40* | 0.496 | Mannose-6-phosphate isomerase, class I |
| CNAG_04879 | *GDB1* | 0.459 | Glycogen debranching enzyme |
| CNAG_06096 |  | 0.472 | Tricarboxylate carrier |
| CNAG_06374 | *MAE1* | 0.267 | Malate dehydrogenase (oxaloacetate-decarboxylating)(NADP) |
| CNAG_06923 | *XFP2* | 0.335 | Xylulose 5-phosphate/fructose 6-phosphate phosphoketolase |
| CNAG_07745 | *MPD1* | 0.186 | Alcohol dehydrogenase, propanol-preferring |
| **DNA replication, cell cycle and differentiation** | | | |
| CNAG_05643 |  | 0.441 | DNA polymerase delta subunit 4 |
| **Integral component of membrane** | | | |
| CNAG_00453 |  | 0.363 | Mitochondrial protein |
| CNAG_01043 |  | 0.413 | Hypothetical protein |
| CNAG_01621 |  | 0.226 | Hypothetical protein |
| CNAG_01761 |  | 0.343 | Hypothetical protein |
| CNAG_03333 |  | 0.388 | Cytoplasmic protein |
| CNAG_03566 |  | 0.313 | Hypothetical protein |
| CNAG_04103 |  | 0.463 | Hypothetical protein |
| CNAG_04658 |  | 0.497 | Hypothetical protein |
| CNAG_05229 |  | 0.453 | Stomatin family protein |
| **Lipid metabolic process** | | | |
| CNAG_00519 | *ERG3* | 0.305 | Lathosterol oxidase |
| CNAG_00834 | *CHO1* | 0.495 | Phosphatidylserine decarboxylase |
| CNAG_01044 | *YPC1* | 0.364 | Alkaline phytoceramidase |
| CNAG_01737 | *ERG25* | 0.436 | C-4 Methyl sterol oxidase, putative |
| CNAG_02553 |  | 0.453 | Short-chain dehydrogenase |
| CNAG_02577 | *SCS7* | 0.43 | Sphingolipid alpha-hydroxylase |
| CNAG_02751 |  | 0.367 | Short-chain dehydrogenase |
| CNAG_04687 | *OLE1* | 0.481 | Stearoyl-CoA desaturase (delta-9 desaturase) |
| CNAG_05496 |  | 0.482 | 2-Isopropylmalate synthase |
| **Oxidoreductase activity** | | | |
| CNAG_00115 |  | 0.498 | Chlorophyll synthesis pathway protein |
| CNAG_01102 |  | 0.487 | Oxidoreductase |
| CNAG_01464 | *FHB1* | 0.212 | Flavohemoglobin |
| CNAG_01540 |  | 0.459 | Dehydrogenase |
| CNAG_01558 |  | 0.438 | Chlorophyll synthesis pathway protein BchC |
| CNAG_01577 |  | 0.416 | Glutamate dehydrogenase (NADP) |
| CNAG_01947 |  | 0.491 | 2,4-Dienoyl-CoA reductase |
| CNAG_04471 |  | 0.496 | FAD dependent oxidoreductase |
| CNAG_05842 | *ERG110* | 0.488 | Cytochrome P450 |
| **Transporter activity** | | | |
| CNAG_00539 |  | 0.42 | Membrane transporter |
| CNAG_01118 | *AAP3* | 0.35 | AAT family amino acid transporter |
| CNAG_01925 |  | 0.338 | Hypothetical protein |
| CNAG_02527 |  | 0.488 | Multidrug transporter |
| CNAG_03051 |  | 0.309 | Polyamine transporter |
| CNAG_06204 |  | 0.217 | High-affinity nicotinic acid transporter |
| **Protein catabolic process** | | | |
| CNAG_03128 |  | 0.485 | Gamma-glutamyltransferase |
| CNAG_04656 |  | 0.394 | Arginyl-tRNA-protein transferase |
| **ATP binding** | | | |
| CNAG_04310 |  | 0.498 | Hypothetical protein |
| CNAG_07347 | *HSP104* | 0.477 | Heat shock protein |
| **Chaperone binding** | | | |
| CNAG_02691 |  | 0.387 | Hypothetical protein |
| CNAG_02701 |  | 0.347 | Hypothetical protein |
| **Iron ion homeostasis** | | | |
| CNAG_00315 |  | 0.445 | HHE domain-containing protein |
| **Mitochondrial intermembrane space** | | | |
| CNAG_00929 | *MIX23* | 0.496 | Intermembrane space protein |

*Genes are grouped based on their Gene Ontology (GO) term or predicted biological functions
