## Supplementary material for "Evolutionary unique *N*-glycan-dependent protein quality control system plays pivotal roles in cellular fitness and extracellular vesicle transport in *Cryptococcus neoformans*": Table S4

**Supplementary Table S4. Representative extracellular vesicle (EV)-associated proteins commonly detected in this study and previously reported *Cryptococcus neoformans* EV proteome data sets***

| **Locus tag (CNAG)** | **Protein name** | **Description** | **EV** | **Cell lysate** |
| --- | --- | --- | --- | --- |
| **Enzymes involved in cell surface remodeling** | | | |  |
| CNAG_02225 | Exg104 | Glucan β1,3-glucosidase | 5.031 | 5.532 |
| CNAG_03120 | Ags1 | α1,3-glucan synthase | NS | NS |
| CNAG_02850 | Agn1 | Glucan endo-α1,3-glucosidase | 3.888 | ND |
| CNAG_04861 | Vep8 | Transglucosylase SLT domain- containing protein | 3.898 | ND |
| CNAG_05799 | Cda1 | Chitin deacetylase 1 | 3.922 | NS |
| CNAG_01230 | Cda2 | Chitin deacetylase 2 (MP98) | NS | 2.133 |
| CNAG_01239 | Cda3 | Chitin deacetylase 3 (MP84) | 2.581 | NS |
| CNAG_06501 | Gas1 | β1,3-glucanosyltransferase | NS | NS |
| CNAG_05138 | Spr1 | Glucan β1,3-glucosidase | 0.340 | NS |
| CNAG_01854 | Hep2 | Heparinase II/III family | NS | NS |
| CNAG_05411 | Lpi9 | Endoglucanase | NS | NS |
| CNAG_04969 | Ugd1 | UDP-glucose 6-dehydrogenase | NS | NS |
| CNAG_02748 | Ugp1 | UTP-glucose-1-phosphate uridylyltransferase | NS | NS |
| **Carbohydrate metabolic process** | | | |  |
| CNAG_02189 | Amy1 | α-Amylase | 2.111 | NS |
| CNAG_05264 | AmyA | α-Amylase | 2.047 | ND |
| CNAG_03358 |  | Phosphoglycerate kinase | 0.136 | 0.568 |
| CNAG_03266 | Mdh1 | Malate dehydrogenase | 0.297 | NS |
| CNAG_06699 | Gbp1 | Glyceraldehyde-3-phosphate dehydrogenase | 0.307 | NS |
| CNAG_00061 |  | Citrate synthase | NS | NS |
| CNAG_05907 |  | Pyruvate carboxylase | 0.482 | NS |
| CNAG_03245 |  | Glucose-6-phosphate 1-dehydrogenase | 0.193 | NS |
| CNAG_03916 |  | Glucose-6-phosphate isomerase | 0.195 | NS |
| CNAG_03322 | Usx1 | UDP-glucuronic acid decarboxylase 1 | 0.252 | NS |
| **DNA replication, cell cycle and differentiation** | | | |  |
| CNAG_02338 | Gis2 | Cellular nucleic acid-binding protein | NS | 2.140 |
| CNAG_06747 |  | Histone H2A | NS | NS |
| CNAG_06746 |  | Histone H2B | 0.403 | 0.434 |
| CNAG_01648 |  | Histone H4 | NS | 0.460 |
| CNAG_04577 |  | Nucleoside-diphosphate kinase | NS | 0.298 |
| CNAG_05235 | Bmh2 | 14-3-3 protein | 0.306 | NS |
| CNAG_00483 | Act1 | Actin | 0.407 | NS |
| CNAG_05701 | Tpm1 | Actin lateral binding protein | 0.344 | NS |
| CNAG_03787 |  | Tubulin alpha chain | 0.212 | NS |
| CNAG_01840 |  | Tubulin beta chain | 0.208 | NS |
| **Integral component of membrane** | | | |  |
| CNAG_05277 | Snc2 | Vesicle-associated membrane protein | NS | ND |
| CNAG_04953 | Tsh1 | Pali-domain-containing protein | 5.287 | 0.497 |
| CNAG_05654 | Tsh2 | Pali-domain-containing protein | 2.923 | NS |
| CNAG_06424 | Tsh3 | Claudin family protein | 2.356 | NS |
| CNAG_05615 | Sso1 | Syntaxin 1B/2/3 | 2.065 | NS |
| CNAG_06400 | Pma1 | Plasma membrane ATPase | NS | NS |
| CNAG_02974 |  | Voltage-dependent anion channel protein 2 | NS | NS |
| **Oxidoreductase activity** | | | |  |
| CNAG_01019 | Sod1 | Superoxide dismutase [Cu – Zn] | NS | 2.344 |
| CNAG_04388 | Sod2 | Superoxide dismutase [Mn] | 0.227 | NS |
| CNAG_00407 | Gox1 | Glyoxal oxidase | NS | NS |
| CNAG_02030 | Gox2 | Glyoxal oxidase | NS | NS |
| CNAG_05731 | Gox3 | Glyoxal oxidase | NS | NS |
| CNAG_06241 | Cfo1 | Ferroxidase | NS | 0.379 |
| CNAG_02958 | Cfo2 | Ferroxidase | 0.261 | ND |
| CNAG_06917 | Prx1 | Thiol-specific antioxidant protein | NS | NS |
| CNAG_05847 |  | Thioredoxin reductase | 0.404 | NS |
| **Ribosome biogenesis and RNA processing** | | | | |
| CNAG_04726 |  | 60S ribosomal protein L20 | NS | NS |
| CNAG_00656 |  | Large subunit ribosomal protein L7e | NS | 0.508 |
| CNAG_01049 | Nop10 | H/ACA ribonucleoprotein complex subunit 3 | NS | 0.427 |
| CNAG_04762 |  | Large subunit ribosomal protein L4e | NS | NS |
| CNAG_00034 |  | Large subunit ribosomal protein L9e | NS | NS |
| CNAG_07839 |  | Large subunit ribosomal protein L11 | NS | NS |
| CNAG_06095 |  | Large subunit ribosomal protein L13e | NS | NS |
| CNAG_04799 |  | Large subunit ribosomal protein L14e | NS | NS |
| CNAG_06811 |  | Large ribosomal subunit protein eL22 | 0.328 | NS |
| CNAG_03283 |  | Large subunit ribosomal protein L24e | NS | NS |
| CNAG_00771 |  | Large subunit ribosomal protein L29 | NS | NS |
| CNAG_00232 |  | Large subunit ribosomal protein L30e | NS | NS |
| CNAG_00703 |  | Large subunit ribosomal protein L31e | NS | 0.464 |
| CNAG_06605 |  | Small subunit ribosomal protein S2 | NS | NS |
| CNAG_04114 |  | Small ribosomal subunit protein uS2 | 0.345 | NS |
| CNAG_01170 |  | Small subunit ribosomal protein S17 | 0.464 | NS |
| CNAG_01628 |  | Small subunit ribosomal protein S20 | 0.404 | NS |
| **Protein folding and proteolysis** | | | |  |
| CNAG_02500 | Cne1 | Calnexin | 3.841 | NS |
| CNAG_03682 | Frr1 | FK506-binding protein | NS | NS |
| CNAG_03891 |  | Hsp60-like protein | NS | NS |
| CNAG_06208 | Hsp70 | Heat-shock protein 70 | 0.303 | NS |
| CNAG_01404 |  | Hsp71-like protein | NS | NS |
| CNAG_06150 | Hsp90 | Heat-shock protein 90 | 0.172 | NS |
| CNAG_02801 | Trx1 | Thioredoxin | 0.453 | NS |
| CNAG_01040 |  | Carboxypeptidase | NS | ND |
| **Protein trafficking** | | | | |
| CNAG_05068 |  | GTP-binding protein | 2.074 | NS |
| CNAG_02817 | Sav1/Sec4 | GTP-binding protein | NS | NS |
| **Transporter activity** | | | |  |
| CNAG_00019 | Tim9 | Mitochondrial import inner membrane translocase subunit | 0.478 | 0.359 |
| CNAG_00895 | Zip1 | Solute carrier family 39 (zinc transporter) | 0.054 | 0.140 |
| CNAG_06140 | Acb1 | Long-chain fatty acid transporter | NS | 0.247 |
| CNAG_05788 | Vma10 | V-type proton ATPase | NS | NS |
| CNAG_00730 | Afr1 | ABC multidrug transporter | NS | NS |
| **Other functions** | | | |  |
| CNAG_00588 | Ril2 | Ricin-type beta-trefoil lectin domain-containing protein | 4.539 | ND |
| CNAG_00587 | Ril3 | Ricin-type beta-trefoil lectin domain-containing protein | 6.991 | 2.968 |
| CNAG_02545 | Ipp1 | Inorganic pyrophosphatase | 0.120 | NS |
| CNAG_00417 |  | Elongation factor 1-gamma | 0.229 | NS |
| CNAG_06840 |  | Elongation factor 2 | 0.341 | NS |
| CNAG_03315 | Rho1 | GTP-binding protein | NS | NS |
| CNAG_00776 | Mp88 | Immunoreactive mannoprotein MP88 | 4.786 | NS |

*The information on EV-enriched or EV-associated proteins of *Cryptococcus neoformans* was obtained from the previous papers on *C. neoformans* EV proteomic analysis (Rodrigues et al., 2008; Wolf et al., 2014; Rizzo et al., 2021). Proteins with > 2-fold changes in their abundance are highlighted in red (higher in *ugg1*Δ over WT) or blue (lower in *ugg1*Δ over WT). NS – not significant (p > 0.05) ; ND – not determined
